# Peripheral macrophages causally contribute to disease onset and progression in the *Ndufs4*(KO) model of Leigh syndrome

**DOI:** 10.1101/2023.01.27.525842

**Authors:** Allison R Hanaford, Asheema Khanna, Katerina James, Yihan Chen, Michael Mulholland, Bernhard Kayser, Vivian Truong, Margaret Sedensky, Phil Morgan, Vandana Kalia, Nathan Baerchst, Surojit Sarkar, Simon C Johnson

## Abstract

Subacute necrotizing encephalopathy, or Leigh syndrome (LS), is the most common paediatric presentation of genetic mitochondrial disease. LS is a multi-system disorder with severe neurologic, metabolic, and musculoskeletal symptoms. The presence of progressive, symmetric, necrotizing lesions in the brainstem are a defining feature of the disease, and the major cause of morbidity and mortality, but the mechanisms underlying their pathogenesis have been elusive. Recently, we demonstrated that high-dose pexidartinib, a CSF1R inhibitor, prevents LS CNS lesions and systemic disease in the *Ndufs4*(-/-) mouse model of LS. While the dose-response in this study implicated peripheral immune cells, the immune populations involved have not yet been elucidated. Here, we used a targeted genetic tool, deletion of the colony stimulating factor 1 receptor (CSF1R) macrophage super-enhancer FIRE (*Csf1r*ΔFIRE), to specifically deplete microglia and define the role of microglia in the pathogenesis of LS. Homozygosity for the *Csf1r*ΔFIRE allele ablates microglia in both control and *Ndufs4*(-/-) animals, but onset of CNS lesions and sequalae in the *Ndufs4*(-/-), including mortality, are only marginally impacted by microglia depletion. The overall development of necrotizing CNS lesions is not altered, though microglia remain absent. Finally, histologic analysis of brainstem lesions provides direct evidence of a causal role for peripheral macrophages in the characteristic CNS lesions. These data demonstrate that peripheral macrophages play a key role in the pathogenesis disease in the *Ndufs4*(-/-) model.

## Introduction

Subacute necrotizing encephalomyelopathy, Leigh syndrome (LS), is the most common paediatric presentation of genetic mitochondrial disease [13, 14]. LS neuropathology is characterized by symmetric, bilateral, neurodegenerative lesions which can appear in the basal ganglia, brainstem, and cerebellum [8, 13]. Patients are often born without evidence of disease, and symptoms typically first present in early childhood with progressive psychomotor regression and symptoms related to brainstem pathology (e.g. respiratory issues, dysphagia) [13]. LS is clinically defined and genetically heterogeneous disease, complicating diagnosis and study. To date, LS has been causally linked to more than 75 distinct genes [14]. These include genes encoded by both the nuclear and mitochondrial genomes, coding for proteins involved in electron transport chain (ETC) or ETC complex assembly factors, mitochondrial transcription/translation factors, and enzymes involved in thiamine metabolism [14].

One causal gene for LS in humans is NDUFS4, coding for a structural/assembly component of ETC complex I. Homozygous deletion of murine *Ndufs4* results in a LS phenotype in mice, and the *Ndufs4*(-/-) mouse is considered the premier animal model of mitochondrial disease. *Ndufs4*(-/-) mice present with a multi-system disorder similar to human patients which includes necrotic lesions in the brainstem, neurological and metabolic dysfunction, and early mortality [12, 25]. Neuroinflammation and lesions also develop in cerebellum. CNS lesions in the brainstem, olfactory bulb, and cerebellum of *Ndufs4*(-/-) mice are characterized by astrocytosis and microgliosis. Advanced brainstem lesions are well demarcated, developing into densely packed IBA1(+) phagocytic cells with astrocyte accumulation (typically observed via staining with GFAP) surrounding the central lesion. The olfactory bulb and cerebellar neuroinflammation also display increased IBA1(+) cell and astrocytosis.

We have recently demonstrated that disease in the *Ndufs4*(-/-) is causally mediated by immune cells, and that treatment with the CSF1R (Colony Stimulating Factor 1 Receptor) inhibitor pexidartinib/PLX3397 can provide potent therapeutic benefits [24]. CSF1R, regulates the differentiation, function, and survival of macrophages, including microglia [23]. High dose pexidartinib depletes both CNS and peripheral macrophages, but the effect is relatively microglia-specific when administered at low dose, and low dose pexidartinib has been reproducibly used as an experimental tool for microglia depletion [2, 17].

Phagocytic cell accumulation in LS lesions has been interpreted as microgliosis, with no consideration of a role for peripheral cells [16, 19]. However, histologic methods and markers used in prior studies have been non-specific, including the pan-macrophage marker IBA1 and morphologic analysis [9, 19]. In our recent work, we found that low doses of the pexidartinib (sufficient to deplete microglia but not peripheral macrophages) fail to substantially alter disease course while high doses prevent disease. Moreover, staining of lesions using a TMEM119-GFP reporter revealed that a substantial fraction of IBA1(+) cells in CNS lesions do not express this microglia-specific marker [24, 30]. Defining the roles of individual phagocytic cell populations is important for developing immune-targeting strategies into therapies, as CNS microglia and peripheral macrophages have distinct pharmacological properties and targets. In addition, defining the relative roles of peripheral and central macrophages will have significant impact on our basic understanding of the pathogenesis of this mitochondrial disease.

To determine the role of microglia in the pathobiology of LS, here we utilize a genetic model for microglia depletion, the *Csf1r*ΔFIRE mouse [21]. These animals carry deletion of an *fms-*intronic regulator element (FIRE) in the *Csf1r* gene, an element which regulates expression of *Csf1r* in microglia [7, 21]. Mice homozygous for the *Csf1r*ΔFIRE allele (here abbreviated *Csf1r*(fr/fr) with *Csf1r*(wt/fr) and *Csf1r*(wt/wt) denoting heterozygous and wildtype for this allele, respectively), lack microglia and some tissue macrophage populations, but have intact circulating macrophages [21]. Here, we report the impact of genetic depletion of microglia on disease and survival in the *Ndufs4*(-/-). In addition, we directly assess the presence of microglia and peripheral macrophages to LS CNS lesions. Our findings support a model whereby peripheral macrophages are a primary causal cell type in the pathogenesis of LS CNS lesions.

## Results

### Homozygosity for the Csf1rΔFIRE allele results in depletion of microglia in control and Ndufs4(KO) mice

To confirm the impact of the *Csf1r*ΔFIRE allele on microglia populations in our mouse model, we generated an *Ndufs4*(+/-)/*Csf1r*(wt/fr) colony (*Csf1r*ΔFIRE mice in the C57Bl/6 background generously provided by the Blurton-Jones laboratory at UC Irvine, derived from animals provided by the Pridans laboratory at University of Edinburgh, see **Methods** for breeding and backcrossing details). Immunofluorescent staining for IBA1 in controls animals for the *Ndufs4* gene confirmed the impact of the *Csf1r*ΔFIRE deletion on microglia: *Ndufs4*(control)*/Csf1r*(fr/fr) samples were devoid of IBA1 positive cells in the brain, consistent with prior reports of the impact of the *Csf1r*^ΔFIRE^ allele (**Figure 1 A-G**, see **Methods** for genotype details) [21]. Heterozygosity for the *Csf1r*^ΔFIRE^ allele in Ndufs4(control)/Csf1r(wt/fr) mice did not significantly impact microglia numbers compared to Ndufs4(control)/*Csf1r*(wt/wt) animals (**Figure 1 B-G)**.

**Figure 1.**
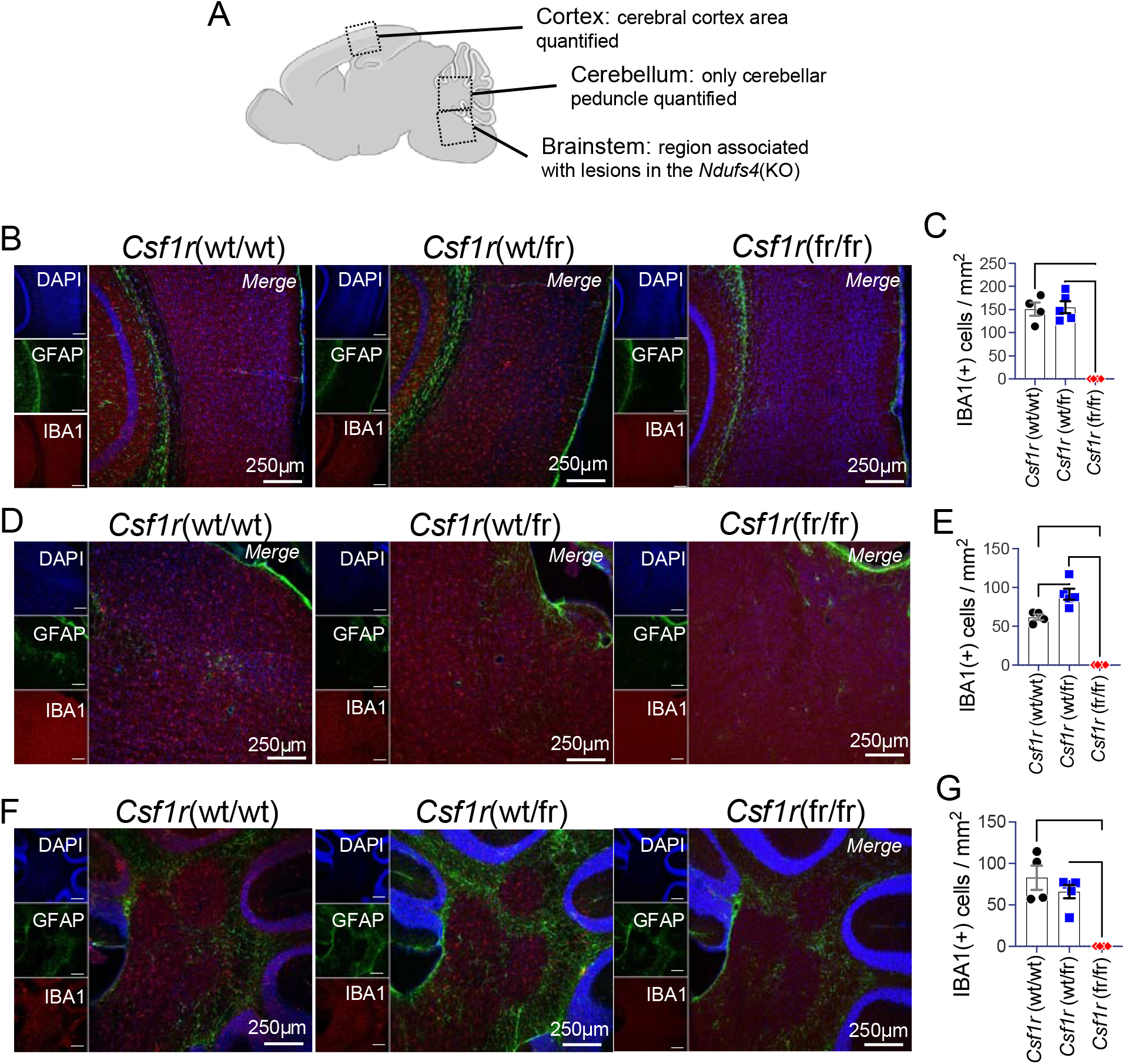
Depletion of microglia in *Csf1r*(fr/fr)mice. (A) Diagram of brain regions imaged and analyzed. In cortex, microglia numbers were only quantified in the cerebral cortex area. In cerebellum, microglia were quantified in the peduncle. In the brainstem, microglia were quantified within the brainstem region where CNS lesions develop in the *Ndufs4*(-/-) model, adjacent to the fourth ventricle and inferior to the cerebellum. Only *Ndufs4* controls are assessed in this figure (see **Methods** for genotype details, **Figures 3-4**). (B) Representative images of cortex from *Csf1r*(wt/wt), *Csf1r*(wt/fr), and *Csf1r*(fr/fr), *Ndufs4*(control), mice. (C) Quantification of microglia numbers within the cerebral cortex. ****p<0.0001 by one-way ANOVA. (D) Representative images of the LS CNS lesion associated region of the brainstem in *Csf1r*(wt/wt), *Csf1r*(wt/fr), and *Csf1r*(fr/fr), *Ndufs4*(control), mice. (E) Quantification of microglia numbers within the lesion-associated brainstem region. ****p<0.0001 by one-way ANOVA. (F) Representative images of the cerebellar peduncle in *Csf1r*(wt/wt), *Csf1r*(wt/fr), and *Csf1r*(fr/fr), *Ndufs4*(control), mice. (G) Quantification of microglia numbers within the cerebellar peduncle. ***p=0.003 by one-way ANOVA. (C, E, G) Error bars are standard error of the mean. Tukey’s multiple comparisons corrected p-values for pairwise comparisons. **p<0.005, ***p<0.0005, ****p<0.00005. n=4 biological replicates per group per brain regions (see **Methods**).

Despite lacking microglia, we observed that, as previously reported, *Csf1r*(fr/fr) mice have no overt phenotype with the exception of an increased incidence hydrocephalus (observed in ∼15% *Csf1r*(fr/fr) pups, versus <1% in C57Bl/6 per Jackson laboratories data). These observations generally agree with prior reports of a hydrocephalus rate of ∼5% in *Csf1r*(fr/fr) animals, and hydrocephaly in *Csf1r* null mice [4, 20]. Hydrocephalus is already increased in many inbred mouse strains, including C57Bl/6, and as standard practice pups with hydrocephaly are euthanized regardless of genotype and not included in experiments (see **Methods**).

### Neuroinflammation in microglia-depleted Ndufs4(KO) animals

We next assessed the impact of the *Csf1r*ΔFIRE allele on neuroinflammation in *Ndufs4*(-/-) mice. Immunofluorescent staining of cerebellum and ventral brainstem demonstrated that while a single copy of the *Csf1r*ΔFIRE allele did not significantly impact microglia numbers, *Ndufs4*(-/-)/*Csf1r*(ft/fr) mice lack IBA1(+) cells in cortex and in brainstem areas not associated with CNS lesions in the *Ndufs4*(-/-) (**Figure 2**).

**Figure 2.**
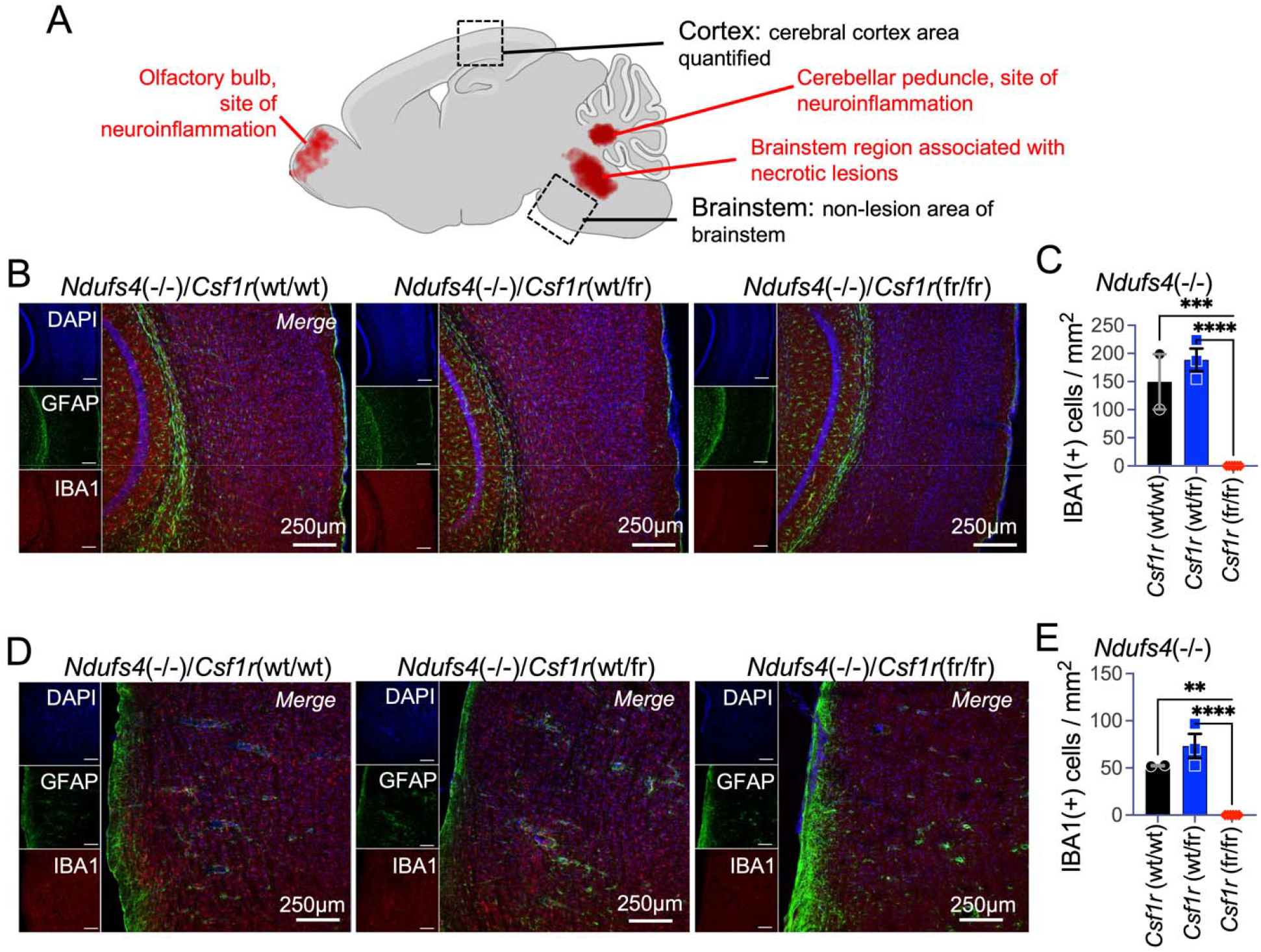
Depletion of microglia in *Ndufs4*(-/-)/*Csf1r*(fr/fr)mice. (A) Diagram of brain region associated with neuroinflammatory lesions in the *Ndufs4*(-/-) model (red highlighted), and the cortex and brainstem regions imaged and analyzed in this figure. (B) Representative images of cortex from *Ndufs4*(-/-)/Csf1r(wt/wt), *Ndufs4*(-/-)/*Csf1r*(wt/fr), and *Ndufs4*(-/-)/*Csf1r*(fr/fr) mice. (C) Quantification of microglia numbers within the cerebral cortex. ****p<0.0001 by one-way ANOVA. (D) Representative images of ventral brainstem region adjacent to areas of lesion formation in *Ndufs4*(-/-)/*Csf1r*(wt/wt), *Ndufs4*(-/-)/*Csf1r*(wt/fr), and *Ndufs4*(-/-)/*Csf1r*(fr/fr) mice. (E) Quantification of microglia number within the lesion-associated brainstem region. ****p<0.0001 by one-way ANOVA. (C, E) Error bars are standard error of the mean. Tukey’s multiple comparisons corrected p-values for pairwise comparisons. **p<0.005, ***p<0.0005, ****p<0.00005. n=2-3 biological replicates per group per brain regions (see **Methods**).

While microglia were absent in regions not associated with lesion formation, necrotizing CNS lesions were not prevented: as in *Ndufs4*(-/-) mice with wildtype *Csf1r, Ndufs4*(-/-)/*Csf1r*(ft/fr) mice at about post-natal day 60 (P60) presented with overt lesions in the brainstem and olfactory bulb comprised of the typical composition of IBA1(+) and GFAP(+) cells (**Figure 3A-G**). Critically, in brain tissue immediately adjacent to the lesions, IBA1(+) cells remain absent (**Figure 2D**; **Figure 3C**, see area adjacent to lesion). The cerebellar peduncle also remained IBA1(+) cell free, possibly uncoupling cerebellar and brainstem lesions, at least at ∼P60 (**Figure 3G**).

**Figure 3.**
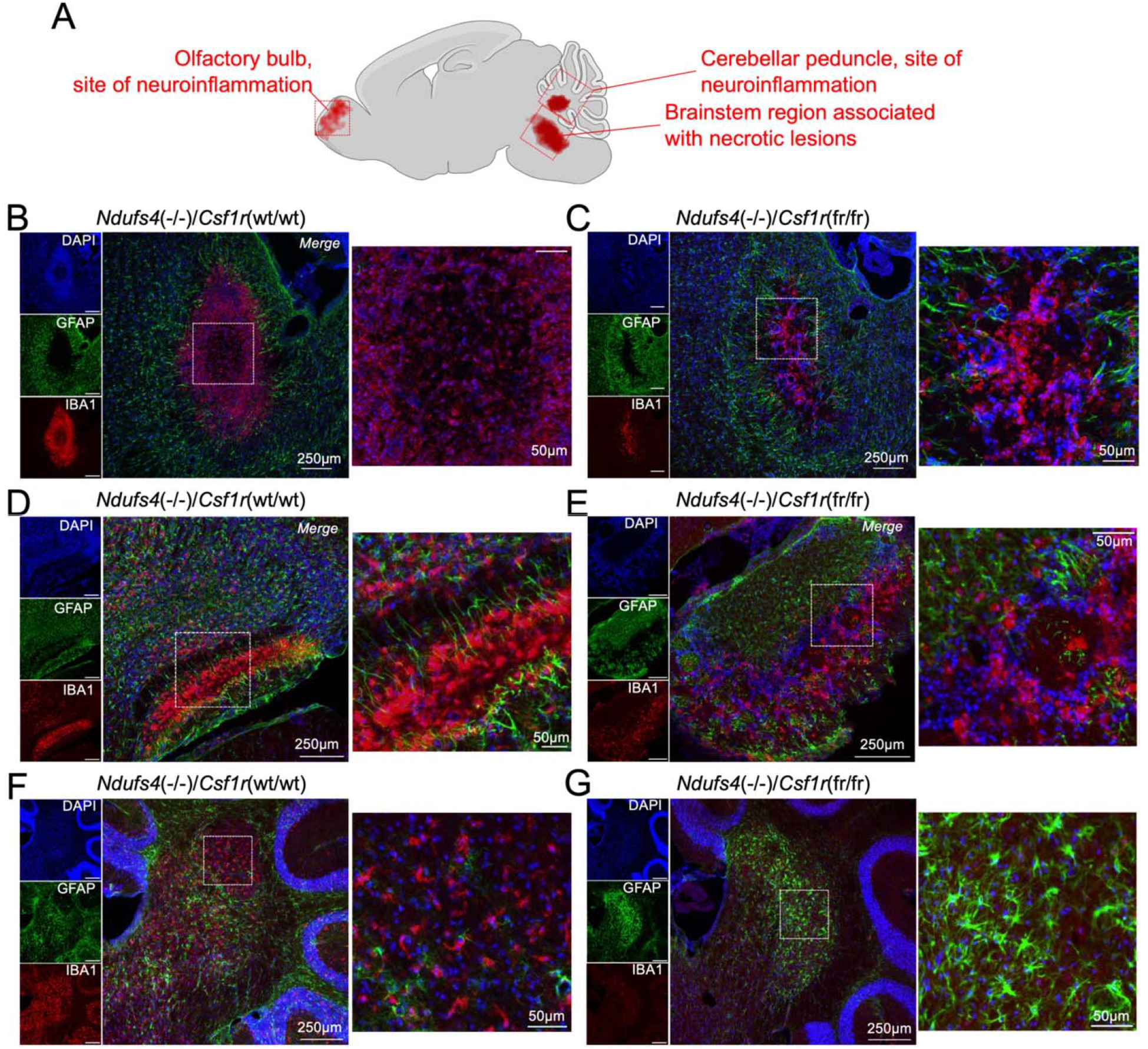
CNS lesions in *Ndufs4*(-/-)/*Csf1r*(fr/fr)mice. (A) Diagram of brain regions associated with neuroinflammatory lesions in the *Ndufs4*(-/-) model and the regions shown in this figure. (B-C) Brainstem CNS lesions in *Ndufs4*(-/-)/*Csf1r*(wt/wt) (B) and *Ndufs4*(-/-)/*Csf1r*(fr/fr) (C) mice. (D-E) Olfactory bulb CNS neuroinflammation in *Ndufs4*(-/-)/*Csf1r*(wt/wt) (D) and *Ndufs4*(-/-)/*Csf1r*(fr/fr) (E) mice. (F-G) Cerebellar peduncle in *Ndufs4*(-/-)/*Csf1r*(wt/wt) (F) and *Ndufs4*(-/-)/*Csf1r*(fr/fr) (G) mice. IBA1(+) cells remain absent in this LS neuroinflammation associated region in *Ndufs4*(-/-)/*Csf1r*(fr/fr) mice. (B-G) All images are representative from staining of at least three biological replicates.

### Impact of Csf1rΔFIRE allele on disease progression and survival in Ndufs4(-/-) mice

To assess the impact of microglia depletion on disease and survival in the *Ndufs4*(-/-) model of LS, we next assess survival of *Ndufs4*(-/-) mice wildtype, heterozygous, or homozygous for the *Csf1*ΔFIRE allele. In the *Ndufs4*(-/-) model, animals lack overt symptoms of disease until around ∼P37 (see [19, 24]). Around this age, animals begin to lose weight (defined here as cachexia), display signs of forelimb clasping and ataxia, develop progressively worsening hypoglycaemia, and show a progressive decline in performance on the rotarod assay for overall endurance and neuromuscular coordination (see **Methods**). Onset and progression of weight loss was not significantly delayed in either *Ndufs4*(-/-)/*Csf1r*(wt/fr) or *Ndufs4*(-/-)/*Csf1r*(fr/fr) animals compared to *Ndufs4*(-/-)/*Csf1r*(wt/wt) mice (**Figure 4A-B**). Moreover, *Ndufs4*(-/-)/*Csf1r*(fr/fr) showed only marginal benefits to symptoms of disease: delays in onset of ataxia and forelimb clasping were statistically significantly, but extremely modest (**Figure 4C**).

**Figure 4.**
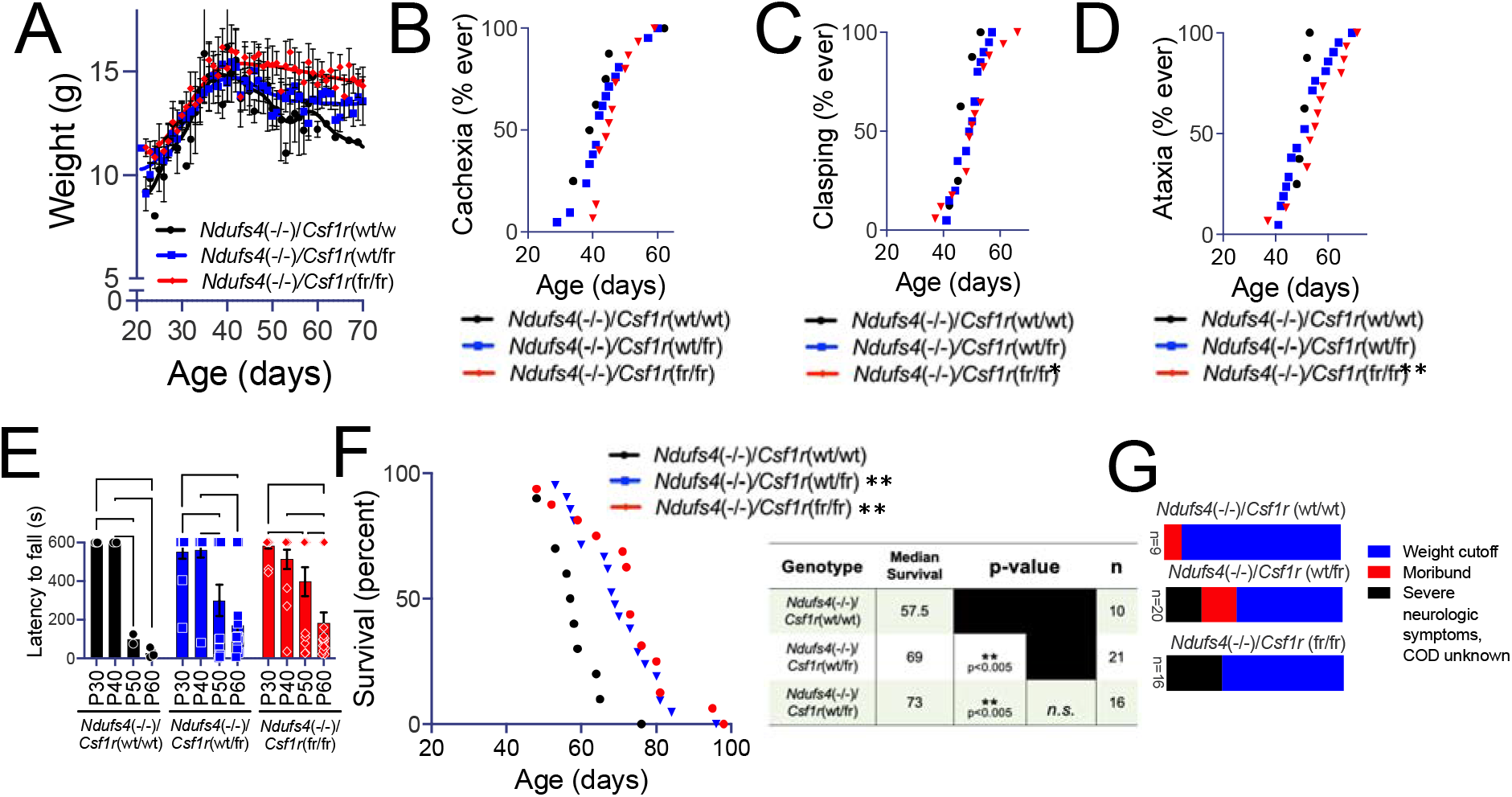
Genetic depletion of microglia provides limited benefits in the *Ndufs4*(-/-) model of LS. (A) Average weights of *Ndufs4*(-/-)/*Csf1r*(wt/wt), *Ndufs4*(-/-)/*Csf1r*(wt/fr), and *Ndufs4*(-/-)/*Csf1r*(fr/fr) mice. Disease onset, including onset of weight loss, occurs around the age of P37 in the *Ndufs4*(-/-) model of LS. Data shown are population averages with standard error of the mean (SEM) and Locally Weighted Scatterplot Smoothing (LOWESS) curves to reveal overall trends. (B) Onset of cachexia, defined as the date when a maximum weight was reached, in *Ndufs4*(-/-)/*Csf1r*(wt/wt), *Ndufs4*(-/-)/*Csf1r*(wt/fr), and *Ndufs4*(-/-)/*Csf1r*(fr/fr) mice. (C) Onset of forelimb clasping in *Ndufs4*(-/-)/*Csf1r*(wt/wt), *Ndufs4*(-/-)/*Csf1r*(wt/fr), and *Ndufs4*(-/-)/*Csf1r*(fr/fr) mice. *p<0.05 by log-rank test. (D) Onset of ataxia in *Ndufs4*(-/-)/*Csf1r*(wt/wt), *Ndufs4*(-/-)/*Csf1r*(wt/fr), and *Ndufs4*(-/-)/*Csf1r*(fr/fr) mice. **p<0.005 by log-rank test. (E) Rotarod performance, as assesed by latency to fall (see **Methods**), in *Ndufs4*(-/-)/*Csf1r*(wt/wt), *Ndufs4*(-/-)/*Csf1r*(wt/fr), and *Ndufs4*(-/-)/*Csf1r*(fr/fr) mice as a function of age. Two-way ANOVA row factor (age) p<0.0005, column factor (genotype) not significant. *p<0.05, **p<0.005, ****p<0.00005 by Tukey’s multiple comparisons corrected t-test performing all age groups within each genotype. (F) Survival of *Ndufs4*(-/-)/*Csf1r*(wt/wt), *Ndufs4*(-/-)/*Csf1r*(wt/fr), and *Ndufs4*(-/-)/*Csf1r*(fr/fr) mice. Median survival and statistical significance, by log-rank test, as shown. (G) Cause of death in *Ndufs4*(-/-)/*Csf1r*(wt/wt), *Ndufs4*(-/-)/*Csf1r*(wt/fr), and *Ndufs4*(-/-)/*Csf1r*(fr/fr) animals.

Median survival was modestly but statistically significantly increased in both *Ndufs4*(-/-)/*Csf1r*(fr/fr) and *Ndufs4*(-/-)/*Csf1r*(wt/fr) animals compared to the *Ndufs4*(-/-)/*Csf1r*(wt/wt) group (**Figure 4F**). In contrast with the benefits of CSF1R inhibition with pexidartinib, respiratory rate defects associated with brainstem lesion progression were not rescued in *Ndufs4*(-/-)/*Csf1r*(fr/fr) and *Ndufs4*(-/-)/*Csf1r*(wt/fr) mice (**Supplemental Figure 1**). Microglia depletion did not alter the primary cause of death in *Ndufs4*(-/-) animals, which was euthanasia due to reaching weight-loss criteria in all *Ndufs4*(KO) cohorts, though a greater fraction of animals died before reaching the 20% weight cut-off (**Figure 4G**, see **Methods**).

### Peripheral macrophages in CNS lesions

Given these findings, we next sought to assess the presence of microglial and peripheral macrophage markers in IBA1(+) cells present in the CNS lesions in the *Ndufs4*(-/-) model. Co-staining of brain slices using antibodies targeting IBA1 and the microglia-specific marker P2YR12 further validated the loss of microglia in cortex (**Figure 5A-B**). In *Ndufs4*(-/-)/*Csf1r*(fr/fr) mice, brainstem lesions were found to be composed of IBA1(+)/P2YR12(-) cells, likely peripheral macrophages (**Figure 5C-D**). In the *Ndufs4*(-/-)/*Csf1r*(wt/wt), IBA1(+) cells in the brainstem lesion show P2YR12 positivity, indicating that microglia make up the bulk of the lesions in animals not carrying the Δ*FIRE* allele, an observation further supported by flow cytometry analysis of brainstem tissue from these animals (**Supplemental Figure 2**). Overall cellularity of the lesions was reduced in *Csf1r*(fr/fr) (as determined by DAPI nuclei density and numbers in the lesion core), and the *Csf1r*(wt/wt) animals show some P2YR12 positivity, supporting the notion that microglia are present in lesions in animals with an intact *CSF1R* allele.

**Figure 5.**
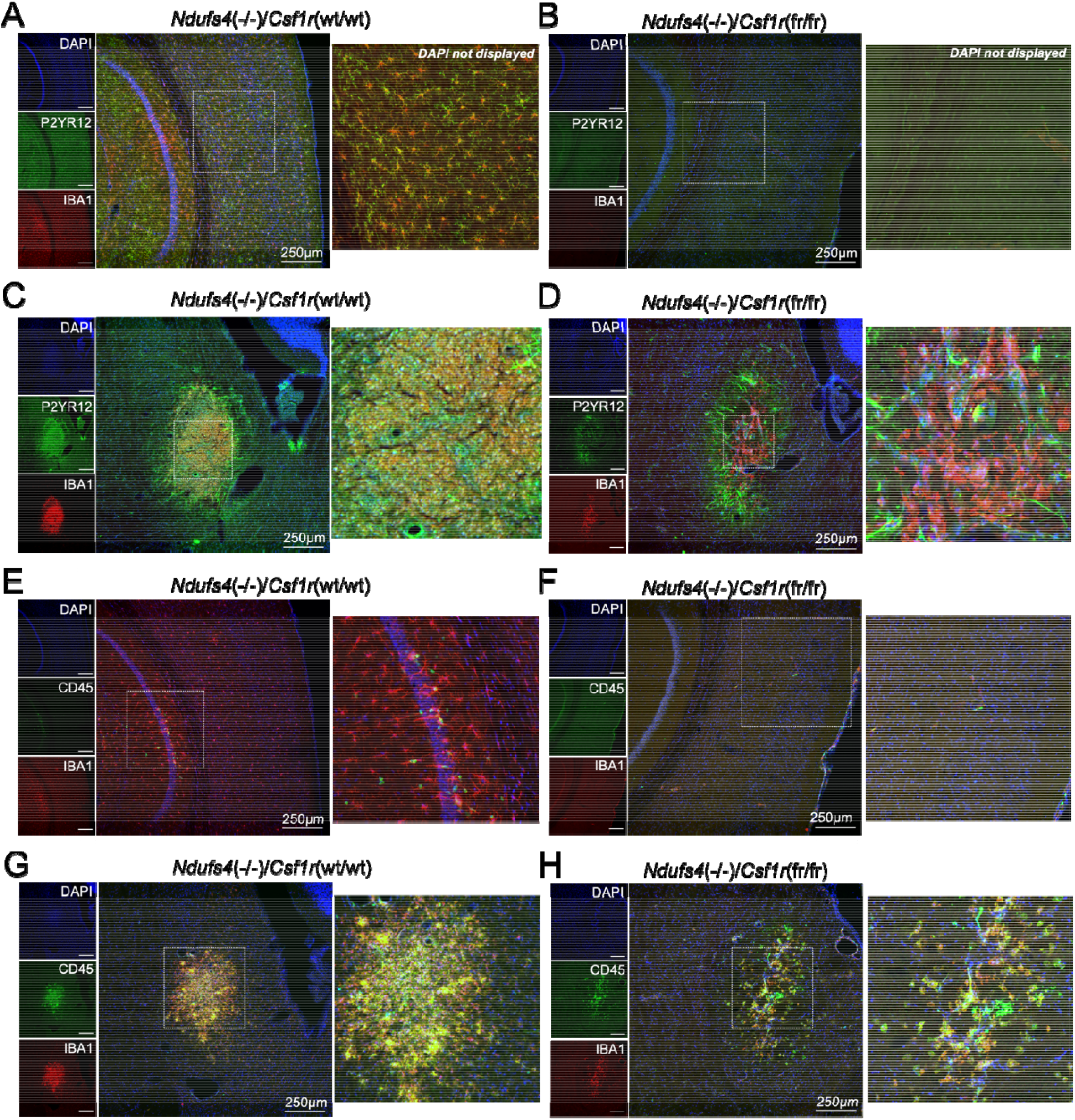
Peripheral macrophages in the CNS lesions characteristic of Leigh syndrome. (A-B) Cortex of *Ndufs4*(-/-)/*Csf1r*(wt/wt) (A) and *Ndufs4*(-/-)/*Csf1r*(fr/fr) (B) mice stained for the pan-macrophage marker IBA1 and the microglia-specific marker P2YR12. Microglia are absent by both IBA1 and P2YR12 staining in the *Ndufs4*(-/-)/*Csf1r*(fr/fr) cortex. (C-D) Brainstem lesions from *Ndufs4*(-/-)/*Csf1r*(wt/wt) (C) and *Ndufs4*(-/-)/*Csf1r*(fr/fr) (D) mice stained for the pan-macrophage marker IBA1 and the microglia-specific marker P2YR12. Cells positive for IBA1, the pan-macrophage marker, are present in both genotypes, while P2YR12 staining is absent in cells in the brainstem lesions of *Ndufs4*(-/-)/*Csf1r*(fr/fr) animals. Note: the P2YR12 staining surrounding the lesion site does not appear to be cellular in origin, and is thought to reflect the presence of aggregated platelets, which are P2YR12 positive. (E-F) Cortex of *Ndufs4*(-/-)/*Csf1r*(wt/wt) (E) and *Ndufs4*(-/-)/*Csf1r*(fr/fr) (F) mice stained for the pan-macrophage marker IBA1 and the peripheral leukocyte marker CD45. Microglia are absent in the *Ndufs4*(-/-)/*Csf1r*(fr/fr) cortex, and microglia (by IBA1 posivitivy and morphology) do not express CD45 A few compact cells with CD45 positivity are present in both genotypes, presumed to be circulating leukocytes. (G-H) Brainstem lesions in *Ndufs4*(-/-)/*Csf1r*(wt/wt) (G) and *Ndufs4*(-/-)/*Csf1r*(fr/fr) (H) mice stained for the pan-macrophage marker IBA1 and the peripheral leukocyte marker CD45. CD45 positive cells are present in both in the *Ndufs4*(-/-)/*Csf1r*(fr/fr) and *Ndufs4*(-/-)/*Csf1r*(wt/wt) lesions, while most or all IBA1 positive cells in the *Ndufs4*(-/-)/*Csf1r*(fr/fr) lesion appear to be positive for the peripheral leukocyte marker CD45. Co-staining of CD45 and IBA1 is indicative of peripheral macrophages.

Similarly, co-staining with IBA1 and the peripheral (haematopoietic origin) immune cell marker CD45 revealed that IBA1(+) cells in *Ndufs4*(-/-)/*Csf1r*(fr/fr) mice are, indeed, peripheral in origin (**Figure 5E-H**). Staining of *Ndufs4*(-/-)/*Csf1r*(wt/wt) mice verified that microglia in cortex are CD45(-), while some CD45(+) are present and show non-microglial morphology (**Figure 5E-F**). In brainstem lesions, *Ndufs4*(-/-)/*Csf1r*(wt/wt) mice both show strong staining for CD45 even when microglia are present, as determined by the presence of IBA1(+)/CD45(-) cells (**Figure 5G**). In *Ndufs4*(-/-)/*Csf1r*(fr/fr) animals, the cellularity of lesions appears reduced as was noted above (determined by overall DAPI nuclei numbers and density), and all or nearly all IBA(+) cells present appear to all be positive for CD45.

Together, these data indicate that peripheral macrophages contribute significantly to the cellular composition of CNS lesions in the *Ndufs4*(-/-) mouse model of LS, and that the absence of microglia reduces cellularity but has only modest effects on disease course.

## Discussion

Here, we report that genetic depletion of microglia only modestly delays disease onset and progression in the *Ndufs4*(-/-) model of LS, contrasting with the previously reported rescue of disease resulting from high dose pexidartinib treatment. Genetic depletion of microglia fails to prevent neuroinflammatory CNS lesion formation, the hallmark feature of LS, and associated sequelae of disease. Furthermore, CNS lesions in both *Ndufs4*(-/-)/*Csf1r*(fr/fr) animals include IBA(+)/CD45(+) cells, consistent with our prior findings that CNS lesions contain TMEM119(-)/IBA1(+) cells. Finally, while microglia appear to account for the majority of lesion cellularity in *Ndufs4*(-/-)/*Csf1r*(wt/wt) animals, lesion formation is not prevented in the absence of microglia and *Ndufs4*(-/-)/*Csf1r*(fr/fr) lesions are characterized by IBA(+)/P2YR12(-) peripheral macrophages. Together, these data demonstrate that peripheral macrophages play a prominent role in the pathogenesis of LS, including in CNS disease, and that the absence of microglia does substantially alter disease course when circulating macrophages are intact.

The findings in this study are consistent with recent work by the Hidalgo laboratory, and our own prior work, demonstrating that low dose pexidartinib, used to eliminate microglia while sparing peripheral macrophages, only modestly alters disease course in the *Ndufs4*(KO) [1, 24]. These data indicate that microglia contribute only partially to the pathogenesis of LS and that targeting microglia alone is insufficient to disrupt disease.

While depletion of microglia is not *sufficient* to prevent disease in the LS model, it remains to be determined whether microglia targeting is *necessary* to disrupt disease progression in the context of pharmacologic targeting of the immune system. In our histologic analysis of *Ndufs4*(-/-) mice with intact microglia we see evidence for both microglia and macrophages in overt lesions. It seems likely that both microglia and macrophages causally contribute. The necessity and sufficiency of targeting peripheral macrophages must be assessed in future studies, perhaps through antibody-mediated depletion of circulating macrophages. It is possible that some other pexidartinib responsive cell population mediates the benefits of that compound. Positive identification of the key causal cell types will require significant future work.

It is notable that the *Csf1r*ΔFIRE allele, even in the heterozygous state, provided a modest benefit to disease given that the number of microglia is not measurably reduced in these animals (**Figure 1**). This finding is consistent with the modest benefits of low dose pexidartinib treatment. While we find that low dose pexidartinib is not effective in robustly altering disease course in the *Ndufs4*(-/-), these observations suggest that low dose CSF1R inhibitors may provide benefits in forms of mitochondrial disease which involve CSF1R dependent cells but are less severe, or progress less rapidly. For example, low dose pexidartinib may attenuate disease in mitochondrial cerebellar ataxia caused by *Aifm1* deficiency, where microglial activation occurs in the absence of overt CNS lesions [11, 28]. It may be that differential activation of peripheral versus CNS immunity contributes to the distinct clinical presentations of unique forms of genetic mitochondrial disease, the mechanistic underpinnings of which have so far been elusive.

It is also possible that our findings may be relevant to other human diseases which phenotypically mimic the symmetric neuroinflammatory CNS lesions seen in LS. These include progressive supranuclear palsy (PSP) and Wernicke’s encephalopathy, diseases associated with exposure to environmental toxins which inhibit the mitochondrial ETC and thiamine deficiency, respectively; microglia activation appears to be a prominent factor in both cases [2, 27]. CSF1R targeting has recently been shown to benefit multiple complex neurodegenerative disease models including Parkinson’s, multiple sclerosis, and Alzheimer’s [5, 18, 22, 29]. These findings suggest that the pathways we are probing here may be common to a range of diseases where evidence suggests causal roles for both immune activity and mitochondrial dysfunction. Future studies will be required to probe links to our findings in the setting of genetic mitochondrial dysfunction and other forms of neurodegeneration, including age-related neurodegenerative diseases.

Many questions remain to be answered. For example, the nature of the inciting immune activating signal remains unknown. Nevertheless, we believe that there is a high potential for clinical translatability of recent work in this field. Supporting this notion, various case reports have indicated that immune modulatory therapies have altered disease course in LS and other genetic mitochondrial diseases [3, 6, 15, 26] (evidence linking immune function to the pathogenesis of mitochondrial disease is summarized here [6]). The data presented here have important implications for any future efforts to translate immune-targeting strategies into treatments for genetic mitochondrial disease.

## Methods

### Animals

*Csf1r*^ΔFIRE/ΔFIRE^ mice are described in Rojo et al [21] and were generously provided by the Jones laboratory at UC Irvine, originally provided to them by Dr Pridans at the University of Edinburgh. *Ndufs4*(+/-) mice were originally obtained from the Palmiter laboratory at University of Washington, Seattle, Washington USA, but are also available from the Jackson Laboratory (strain, 027058). Strain details are described in Kruse et al [12]. Both the *Ndufs4* and *Csf1r*^ΔFIRE/ΔFIRE^ lines are on the C57/BL6 background. *Ndufs4*(-/-) mice cannot be used for breeding due to their short lifespan. *Ndufs4*(+/-) mice were bred with *Csf1r*^ΔFIRE/ΔFIRE^ mice to produce double heterozygous offspring, which were then crossed to produce *Ndfus4*(-/-)/ *Csf1r*^wt/ΔFIRE^ and *Ndfus4*(-/-)/ *Csf1r*^ΔFIRE/ΔFIRE^ mice.

Mice were weaned at P20-22 days of age. *Ndufs4*(-/-) animals were housed with control littermates for warmth as *Ndufs4*(-/-) mice have low body temperature [12]. Mice were weighed and health assessed a minimum of 3 times a week. Wet food was provided in the bottom of the cage to *Ndufs4*(-/-) mice following symptom onset. Animals were euthanized if they reached a 20% loss of maximum body weight (measured two days consecutively), were immobile, or found moribund. Mice heterozygous for *Ndufs4* have no reported phenotype, so controls consisted of both heterozygous and wild-type *Ndufs4* animals. The *Ndufs4*(Ctrl) and *Ndufs4*(-/-) mice wild type for FIRE (*Csf1*^*wt*/wt^) used in this study came from crosses of *Ndufs4*(+/-)/*Csf1*^wt/ΔFIRE^ mice or from our general Ndufs4 colony. Mice were fed PicoLab Diet 5058 and were on a 12-hour light-dark cycle. All animal experiments followed Seattle Children’s Research Institute guidelines and were approved by the SCRI IACUC.

Clasping, circling, and ataxia were assessed by visual scoring and analyzed as previously described [10]. During disease progression *Ndufs4*(-/-) animals can display intermittent/transient improvement of symptoms. Here we report whether the animal *ever* displayed the symptoms for two or more consecutive days.

The rotarod performance test was performed using a Med Associates ENV-571M single-lane rotarod. A mouse was placed on the rod already rotating at 6 rpm and latency to fall was timed for a maximum of 600 seconds while rotation remained constant. For each mouse, three trials were performed with a minimum of 10 minutes between each trial. The best of three trials was reported.

### Respiratory function

Breathing parameters were recorded from alert, unrestrained, ∼P60 (+/-2 days) mice using whole-body plethysmography. Paired 300 ml recording and reference chambers were continuously ventilated (150ml/min) normal air (79%N_2_/21%O_2_). Pressure differences between the recording and reference chambers were measured (Buxco) and digitized (Axon Instruments) to visualize respiratory pattern. Mice were allowed to acclimate to the chambers for 30-40min prior to acquisition of 35min of respiratory activity in normal air. Respiratory response to hypercapnia was then tested by ventilating the chambers with hypercapnic gas for 15min. Respiratory frequency and breath-to-breath irregularity scores 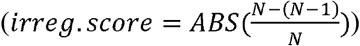 of frequency and amplitude (peak inspiratory airflow) were quantified during periods of resting breathing (pClamp10 software).

### Brain collection, immune-staining, and imaging

Tissue was collected for *Ndufs4*(Ctrl) mice between P60-P66 and for *Ndufs4*(-/-) mice when euthanasia criteria were met. Mice were fixed-perfused with 20mL of PBS followed by 20mL of 4% PFA. Immediately following perfusion, the brain was removed and placed in 4% PFA for a minimum of 24h at 4C. The brain was then moved to cryoprotection solution (30% sucrose, 1% DMSO, 100uM glycine, 1x PBS) for a minimum of 48h at 4C. Cryopreserved tissue was frozen in OCT in cryoblocks on dry ice and stored at -80C until cryosectioning. Cryoblocks were sectioned at 50μm thickness using a Leica CM30505 cyrostat set at -20C. Slices were immediately placed in 1xPBS with DAPI at 4C for a maximum of 24h before mounting on slides. Prior to mounting, slices were examined for the presence lesions using a fluorescent microscope. Slices with lesions were mounted on superfrost plus slides. Four to five slices were mounted on each slide. For control animals, similar anatomic sections were chosen for mounting. Sections from the center of the lesion were chosen for staining and are shown here.

Mounted sections were baked at 37C in a white-light LED illuminated incubator for 24h before storing at -80C until staining. For immunofluorescent staining, slides were incubated for 24h at 60C in pH 6.0 citrate antigen retrieval buffer. Following antigen retrieval, slides were washed for 5 minutes in PBS on ice before being incubated for 1h on ice in 1mg/mL sodium borohydride in PBS. Slides were then washed in PBS with 10mM glycine for 5 minutes before being placed in a 0.5mg/mL sudan black in 70% ethanol and incubated overnight with gentle stirring. Slides were then washed for 5 minutes 3x in PBS. Excess moisture was wiped away and a hybriwell sealing sticker (Grace Bio-Labs, GBL612202) applied to each slide over the tissue slices. Slides were incubated for 30 minutes at room temperature in blocking/permeabilization solution (1mM digitonin, 10% rabbit serum, 0.5% tween in PBS). Following blocking, the hybriwell sealing stickers were removed and blocking solution shaken off the slides. Excess moisture was wiped away and new hybriwells placed. The slides were incubated for 24h at 4C in antibodies and DAPI (1ug/mL) diluted in the blocking solution. The following fluorescently conjugated primary antibodies were used: Anti-IBA1 (1:150, Wa-/-013-26471), anti-GFAP (1:300, clone GA5, Invitrogen 14-9892-82). After incubation, slides were washed 4x 5 min in PBS. ProLong Gold Antifade was used as mounting media and coverslips sealed with clear nail polish. Slides were stored in an opaque slide box at 4C. The following unconjugated primary antibody was used: P2RY12 (1:100, clone S16D007D, Biolegend, 848002. Unconjugated primary and conjugated primary were both included in the initial 24h incubation, followed by 4x 5 min washes in PBS. Slides were then incubated in conjugated secondary antibody (1:500, anti-rat IgG AF-488 conjugate, Cell Signalling Technologies, #4416) and DAPI (1ug/mL) for 2h at room temperature. Slides were then washed and cover slipped as described above.

Imaging was performed on a Zeiss LSM710 confocal microscope. Images were collected using a 10X objective with a 0.6X optical zoom. Channels were set to an optical thickness of 15um or 25nm. 15um images were used for quantifying IBA1(+) cells. 25um images are shown in Figures 3 and 5. DAPI was excited with a 405nm laser, Alexa5488 with a 488nm laser, and fluorochrome 635 (IBA1, Wako) at 633nm. Images were taken with the same laser intensity and collection filter settings. Gain/brightness/contrast were arbitrarily adjusted for visualization to make cell morphology clearly visible (intensities not compared).

IBA1(+) cells were counted using ImageJ. Brightness and contrast were adjusted in ImageJ to enhance visualization of cell morphology. Cells were counted based on IBA1 positivity, not intensity, so this does not impact the results. Image areas of roughly the same size ad anatomical location were chosen and all cells in that region were counted and the number of cells per mm^2^ calculated. Only one image from each mouse was counted.

### Statistical analysis

All statistical analyses were performed using GraphPad Prism. Error bars represent standard error of the mean (SEM) and *P*<0.05 is considered statistically significant.

### Scientific rigor

*Sex*-Approximately equal numbers of male and female animals were used in each experiment. No significant sex differences have been reported in *Ndfus4(-/-)* or *Csf1r*^ΔFIRE/ΔFIRE^ mice and none were observed here.

### Exclusion criteria

Animals euthanized prior to the age of disease onset in the *Ndufs4*(-/-) were excluded from study. Our criteria for early life exclusion includes severe weaning stress (significant weight loss or spontaneous mortality before P30), runts (defined as ≤ 5 g body weight at weaning age), or those or born with health issues unrelated to the *Ndufs4*(-/-) phenotype (such as hydrocephalus). These criteria are applied to all genotypes as part of our standard animal care.

**Figure S1.**
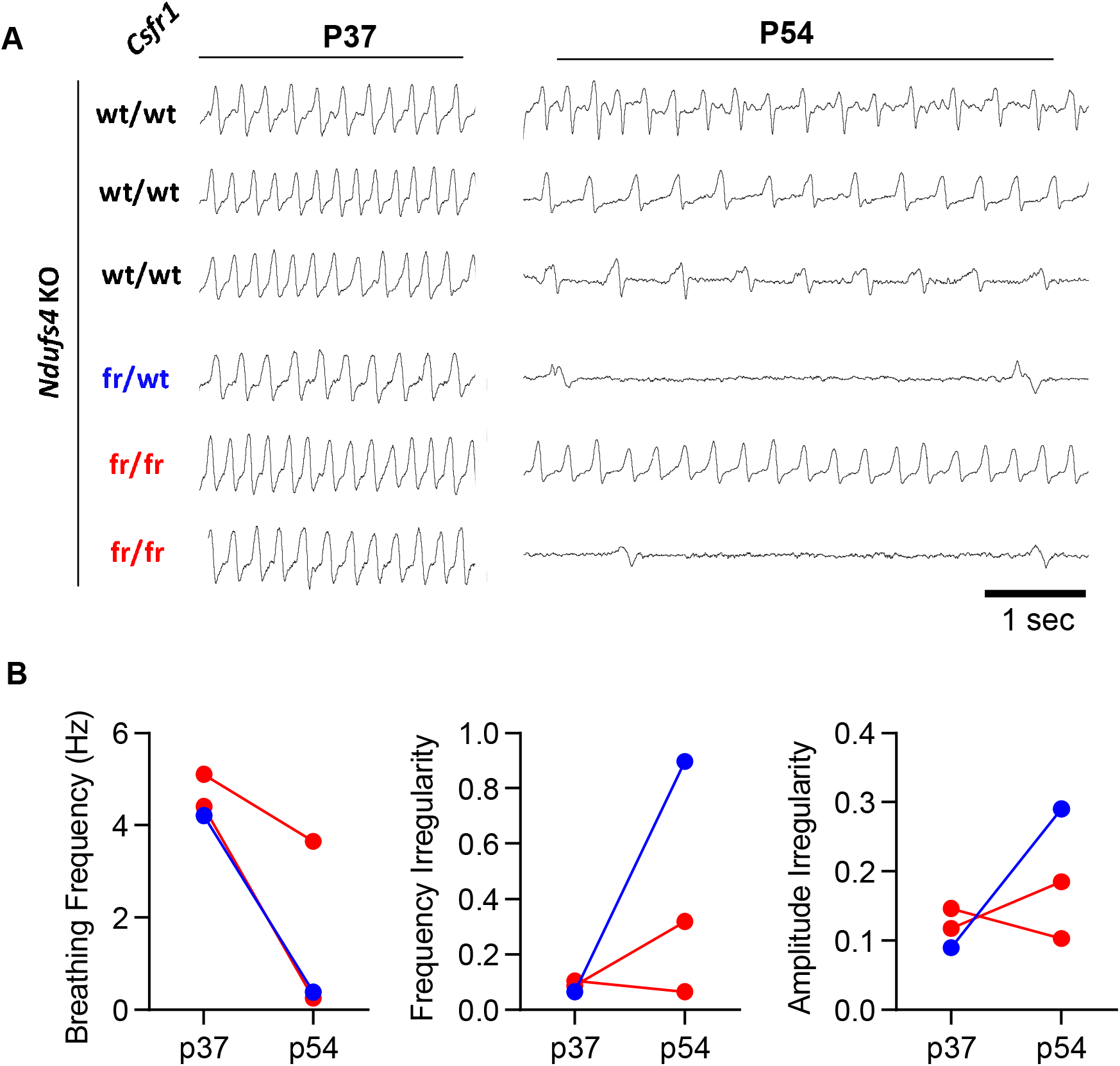
Respiratory defects are not rescued by microglial depletion in *Ndufs4*(-/-) mice. (A) Representative respiratory tracings by plethysmography in *Ndufs4*(-/-) mice wildtype, heterozygous, or homozygous for the *Csf1r*ΔFIRE allele. (B) Plots of breathing rate frequency, rate of frequency irregularities, and rate of amplitude irregularities in *Ndufs4*(-/-) mice wildtype, heterozygous, or homozygous for the *Csf1r*ΔFIRE allele. Contrasting with our previously published results using the CSF1R inhibitor pexidartinib (Stokes et al, 2022), this pilot cohort demonstrated that microglial depletion did not prevent respiratory defects in the *Ndufs4*(-/-) model of Leigh syndrome.

**Figure S2.**
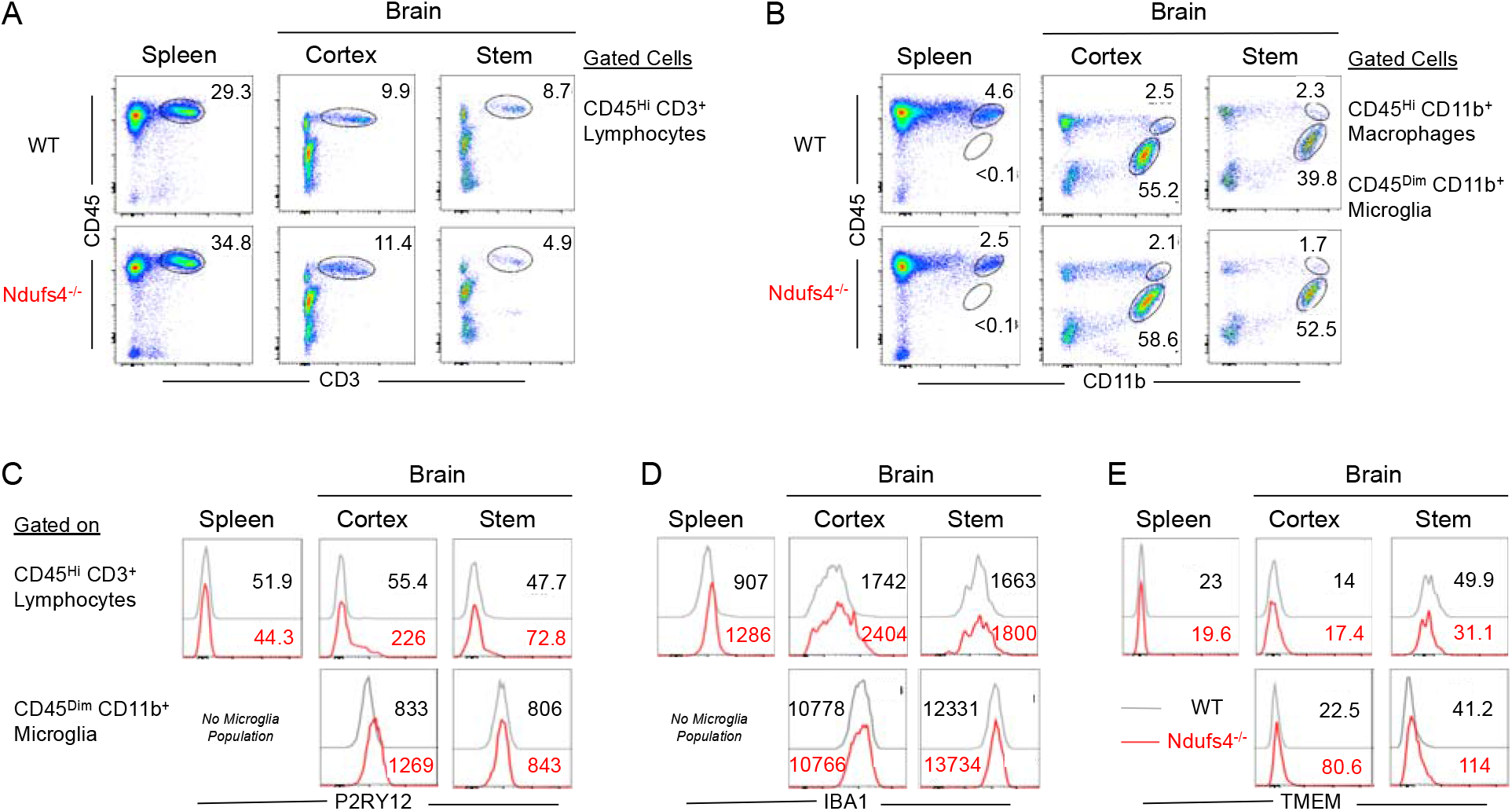
Immunological characterization of hematopoietic lineage cells in spleen, brain cortex and brain stem isolated from wild type and *Ndufs4*(-/-)/*Csf1r*(wt/wt) mice. Spleen, brain cortex and brain stem were isolated from 60 days old wild-type (WT) and diseased *Ndufs4*(-/-)/*Csf1r*(wt/wt) mice. **(A)** Live cells were analyzed for CD45 and CD3 to determine the proportions of CD3+ CD45^Hi^ lymphocytes in indicated tissues. Numbers in flow plots indicate proportions of gated subsets. (**B)** CD45 and CD11b markers were used to distinguish CD45^Hi^ CD11b+ macrophages and CD45^Dim^ CD11b+ microglial cells. Numbers in flow plots indicate proportions of gated subsets. Gated lymphocyte and microglial immune subsets from spleen, cortex and brain stem are presented as histogram plots for expression levels of **(C)** P2RY12, **(D)** IBA1 and **(E)** TMEM in healthy WT and diseased KO mice. Numbers in the histogram plots indicate mean fluorescence intensity (MFI) of expression of indicated markers. RESULTS: Towards characterizing the hematopoietic lineage cell subsets associated with Leigh Syndrome, we conducted a comparative analysis of lymphocytes, macrophages and microglia cells in the diseased brain stems of *Ndufs4*(-/-)/*Csf1r*(wt/wt) and healthy wild-type (WT) mice. Spleens and healthy cortical tissues were also analyzed as controls for immune alterations in tissues without lesions and also as a staining control for microglial cells. Compared to WT mice, *Ndufs4*(-/-)/*Csf1r*(wt/wt) mice showed a modest increase in lymphocyte proportions in spleen and cortex, with a concomitant reduction of lymphocytes in the lesioned brain stems (A). Macrophages showed a universal trend towards reduced proportions in all three tissues (B). In contrast, microglial cells were found to be preferentially enriched in the lesioned brain stems of *Ndufs4*(-/-)/*Csf1r*(wt/wt) mice compared to WT mice. Phenotypically, microglia expressed higher levels of lineage-specific markers P2RY12 (C), as well as the IBA1 marker (D) shared with the macrophage lineage, compared to lymphocytes. Likewise, TMEM19, a microglia specific marker, was expressed at higher levels in microglia compared to lymphocytes (E). Relatively increased expression of P2RY12 (C) and TMEM (E) in the microglia of *Ndufs4*(-/-)/*Csf1r*(wt/wt) compared to WT brain tissues indicates an inflamed phenotype of the microglial cells in the diseased animals. Collectively, these data show global as well as brain stem localized alterations in immune subsets, thus lending further support to immune involvement in Leigh Syndrome. Furthermore, these data indicate that in animals with intact microglia, microglia are the major contributor to CNS lesion mass among phagocytic cell types.

